# The inflammatory skin disease map (ISD map): an interactive computational resource focused on psoriasis and atopic dermatitis molecular mechanisms

**DOI:** 10.1101/2025.02.28.640747

**Authors:** Marcio L. Acencio, Oxana Lopata, Ahmed A. Hemedan, Nick Dand, Stephan Weidinger, Lavinia Paternoster, Sara J. Brown, Joseph Rastrick, Catherine H. Smith, Matthias Hübenthal, Ravi Ramessur, Matladi Ndlovu, Soumyabrata Ghosh, Xinhui Wang, Reinhard Schneider, Venkata Satagopam, Marek Ostaszewski

## Abstract

**Background:** Inflammatory skin diseases (ISD), including atopic dermatitis (AD) and psoriasis (PsO), emerge from a complex network of inter- and intracellular biochemical interactions under the influence of genetic and environmental factors. The complexity of ISD mechanisms hinders translation of research findings into effective treatments and may explain the low remission rates despite the availability of modern targeted therapies.

**Objective:** To model AD- and PsO-associated mechanisms as networks of context-specific molecular interactions, the so-called ISD map, and to check the usefulness of this map as a graphically guided review of AD and PsO mechanisms and as a mechanistic hypothesis-generating platform.

**Methods:** The ISD map was built by assembling mechanistically resolved causal interactions obtained from relevant biomedical literature via manual curation.

**Results:** We demonstrate that the ISD map (https://imi-biomap.elixir-luxembourg.org/) serves as an interactive, graphical review of AD and PsO molecular mechanisms and as a mechanistic hypothesis-generating platform. By analysing the map structure itself or the map integrated with genetics and functional genomics data, we could generate the following mechanistic hypotheses: (i) AD poor response to dupilumab is associated with a potential upregulation of IFNG, IL22, TSLP, IL-17A and IL25 signalling pathways in keratinocytes and/or single nucleotide polymorphisms (SNPs) in genes encoding regulators of IFNG expression in Th1 cells and (ii) PsO resistance to cytokine-induced apoptosis is associated with SNPs in IFNG signalling genes regulating SOCS1 in keratinocytes. Finally, the IL4/IL13 pathway in the AD submap of the ISD map was converted into a probabilistic Boolean model to simulate the effects of IFNG in sensory perception of itching after treatment with dupilumab. Our findings suggest that inhibiting both IFNG and IL4R may improve the therapeutic management of itching.

**Conclusion:** The ISD map provides a significant interactive, computationally accessible resource of molecular knowledge on AD and PsO that can be used to graphically review known AD and PsO mechanisms and generate mechanistic hypotheses.

## INTRODUCTION

Inflammatory skin diseases (ISD) are chronic, non-communicable skin diseases that are associated with genetic predisposition, environmental triggers, metabolic disturbances and dysregulated immune responses. Among the most frequent chronic ISDs are atopic dermatitis (AD), psoriasis (PsO), acne, urticaria, lichen planus, and hidradenitis suppurativa (1). ISDs are an important medical burden because of their high and increasing prevalence and associated comorbidities and psychological impact, as well as diagnostic and therapeutic challenges: there are currently no curative treatments (1).

AD is a common, chronic and relapsing ISD with a global prevalence around 3% (2). AD usually starts in the first 5 years of life, although its onset can occur at any age. AD has an enormous impact on a patient’s quality of life due to intense itch and skin pain (3). Furthermore, there are significant effects on mental health with higher incidence of depression and suicide (4). Similar to AD, PsO is also a common, chronic, and relapsing ISD, with a global prevalence around 3% (5). PsO can occur at any age, but it has two peaks of onset: 20 to 30 years-old and 50 to 60 years-old (6). PsO also can significantly impact the quality of life for the patients (7).

Several molecular mechanisms underlying AD and PsO have been established as targets for innovative therapeutics (8)(9). For example, monoclonal antibodies targeting specific cytokines or their receptors (“biologics”) have emerged as treatment options in AD and PSO. However, full disease control is not always achieved in clinical practice (10)(11), even by the most successful biologic to treat AD, the IL4R inhibitor dupilumab: it induces remission in < 40% of patients when administered as monotherapy (9). This may be due to the fact that IL4R downregulates pathways whose disease-promoting functions can be rescued by alternative pathways that are insensitive to the mechanism of action of the biologic. Therefore, understanding how to target multiple non-redundant pathways and their interconnection with disease-promoting pathways in heterogeneous diseases such as AD and PsO may be translated into a great therapeutic improvement.

As AD and PsO emerge from a complex network of inter- and intracellular biochemical interactions, we expect that the representation of such interactions in the format of a computable network will be useful for the identification of effective therapeutic targets. Such a resource will allow studying the importance of the targets for the disease as measured by high-throughput multi-omics data and observing their position in the network (12). This in turn requires the construction and analysis of a molecular network encompassing PsO- and AD-related causal interactions.

Here, we present the ISD map, a manually-curated network of causal interactions related to AD and PsO (https://imi-biomap.elixir-luxembourg.org) formatted as an interactive diagram of disease mechanisms. We showcase how ISD map can be used as (i) a graphically guided review of PsO and AD molecular pathogenesis and as (ii) a mechanistic hypothesis-generating platform based either on the map structure itself or integration with omics data. The mechanistic hypotheses generated include potential effects of PsO-associated single nucleotide polymorphisms (SNPs) on cytokine-induced apoptosis resistance in psoriatic keratinocytes and potential consequences of AD-associated SNPs on an IFNG-lead poor response to dupilumab in AD. Finally, we describe the results of Boolean network simulations that model the effects of IFNG in sensory perception of itching after treatment with dupilumab in AD. To help users to get acquainted to the navigation and utilization of the ISD map, we provide a quick user guide that can be found in the Supplementary Material.

## METHODS

The ISD map was built via manual curation of the biomedical literature based on the Biomarkers in Atopic Dermatitis and psoriasis (BIOMAP) project experts-suggested key cell types and pathways and potential candidate genes. The ISD map was made publicly available via the MINERVA platform (13) at https://imi-biomap.elixir-luxembourg.org/minerva. More details about the methods can be found in the Supplementary Methods.

## RESULTS

### Scope and structure of the ISD map

AD and PsO are characterised by a chronic inflammatory state that emerges from a complex network of inter- and intracellular biochemical interactions under influence of genetic and environmental factors. The ISD map presents this complex network as a diagrammatic visualisation (Figure 1) using systems biology standards. By using the MINERVA Platform (13)(14), users can explore the map interactively, search for molecules or publications, or visualise omics data by overlaying it over the diagram, contextualising it to related disease mechanisms (please refer to the Supplementary Material for a quick user guide on how to perform these tasks). Contents of the ISD map are annotated using commonly used identifiers, ontologies and controlled vocabularies.

**Figure 1.**
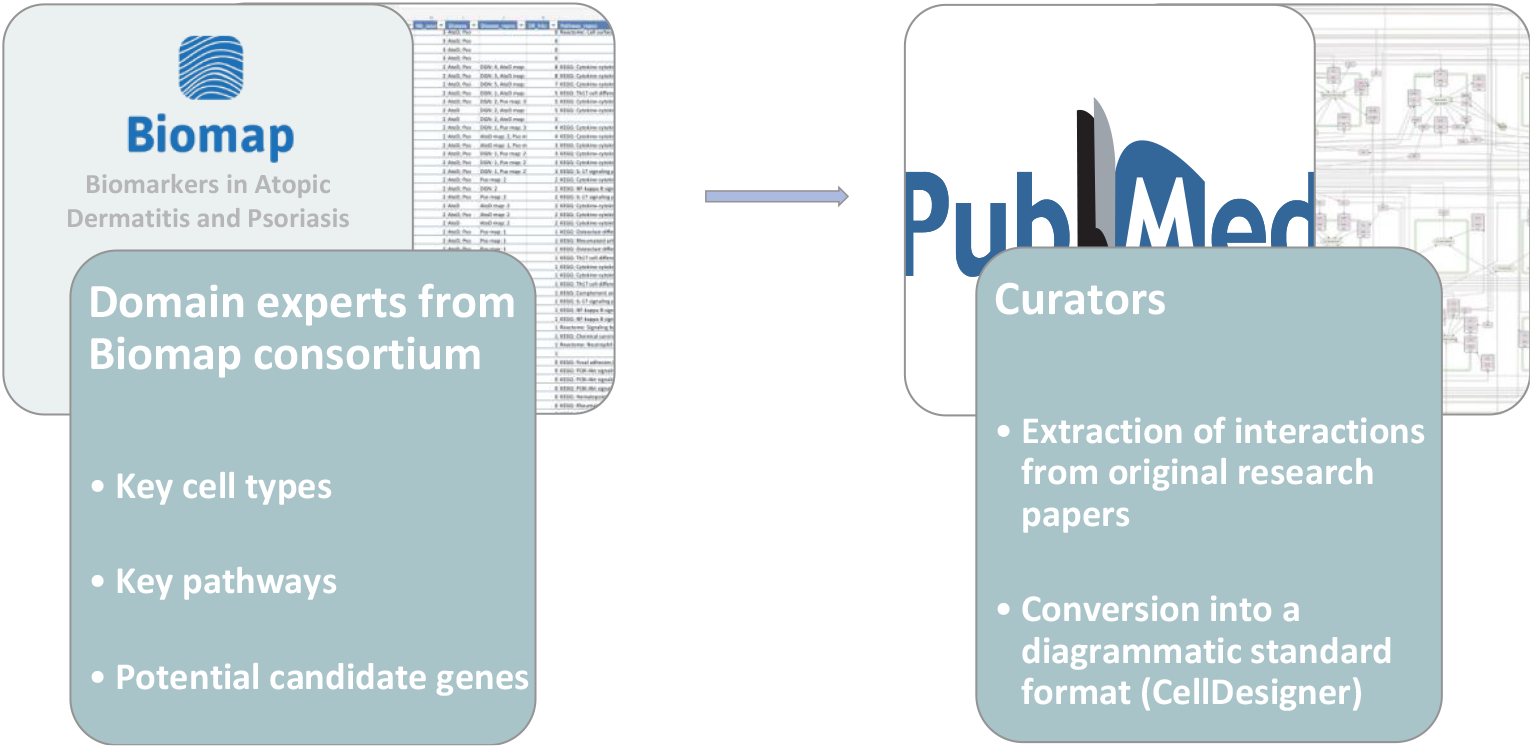
Building the ISD map: Domain experts from the BIOMAP project selected and provided curators with key cell types and pathways and omics-derived potential candidate genes related to atopic dermatitis (AD) and psoriasis (PsO). Curators then extracted relevant causal interactions from original research papers reporting experts’ suggestions and encoded the extracted knowledge into the CellDesigner format. The ISD map was uploaded to the MINERVA platform and then made publicly accessible so that BIOMAP domains experts could review it. After some rounds of reviews and refinement, a final improved version of the map was uploaded to MINERVA.

The ISD map has three layers: (i) a side-by-side AD and PsO overview (Figure 2A), (ii) the intercellular communication view for each disease, and (iii) intracellular pathways (Figure 2B). The side-by-side layer is the entry point for the ISD map and contains the key molecules and cells involved in the onset and progression of AD and PsO (Figure 2A). Intercellular communication views depict how AD and PsO-relevant cell types interact with each other via cytokines, chemokines, metabolites and their receptors. Finally, the intracellular pathways layer show, in some selected cells, details about how the signalling triggered by receptors (Figure 2B) affects the expression of proteins influencing AD- and PsO-related phenotypes.

**Figure 2.**
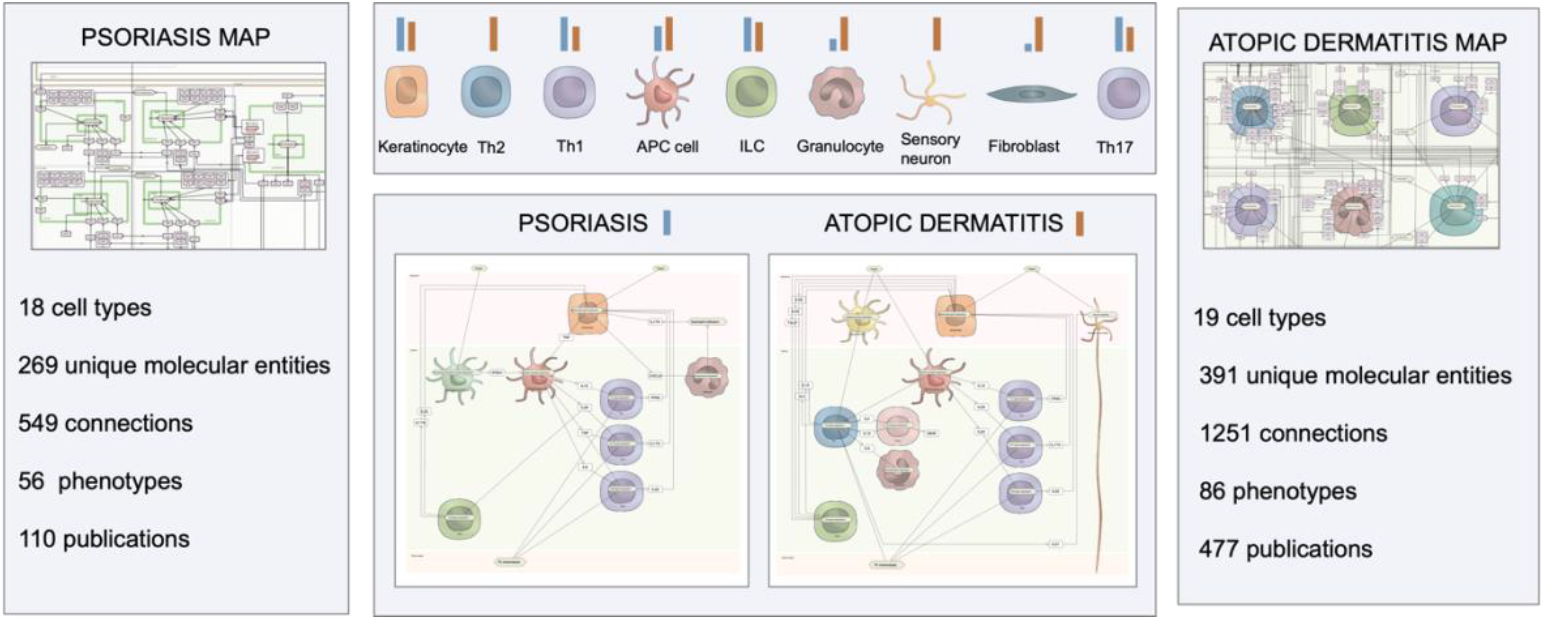
Overall structure of the ISD map: (**A**) The side-by-side overview of the ISD map displays key cells, molecules and cell-cell interactions involved in AD (left panel) and PsO (right panel). (**B**) The intercellular level of ISD map is comprised by different cell types and their relevance (represented as lengths of bars above the respective cells) for AD (orange bars) and PsO (blue bars). The signalling pathways found at the intracellular levels for each disease are listed.

The current versions of AD map (version 02.07.2024) contains 1206 causal interactions, 358 non-redundant biomolecules, 77 processes and 19 different cell types. The current version of PsO map (version 02.07.2024) contains 549 causal interactions, 269 non-redundant biomolecules, 56 processes and 18 different cell types. The causal interactions of both PsO and AD maps are distributed across compartments representing organs and tissues relevant for the disease, such as, for example, epidermis and dermis.

### The ISD map contents: a graphical review of mechanisms

The ISD map was built based on the review of 460 AD- and 110 PsO-related articles and, as such, the exploration of the contents of this map leads naturally to a graphical review of established molecular mechanisms of these diseases. In this section, we summarise the main molecular and cellular aspects of AD and PsO based on the contents of ISD map at the side-by-side overview level (Figure 2A). We provide hyperlinks that allow readers directly access the causal interactions related to a mentioned fact in the ISD map. The detailed description of the mechanisms at the intercellular and intracellular levels – accessible by clicking the buttons “ATOPIC DERMATITIS” and “PSORIASIS” in the side-by-side overview map – can be found in the Supplementary Material.

In AD, initial trigger factors elicit keratinocytes (KCs) to release proinflammatory molecules that activate immune cells such as ILC2 cells (see “ILC2 cells activation by keratinocytes” in the map [at the MINERVA platform](https://imi-biomap.elixir-luxembourg.org/minerva/index.html?id=ISD_map_sidebyside_level&perfectMatch=false&modelId=359&backgroundId=588&x=2470.25&y=1545.75&z=4&overlaysId=1346)). Independently of KCs, some antigens can also be involved in the onset of AD by directly activating Langerhans cells (LCs) and conventional dendritic cell (cDCs). In response to these antigens, LCs and cDCs drive the differentiation of naïve T cells into specific T helper (Th) cells (see “Langerhans and dendritic cells-driven T cell polarization” in the map [at the MINERVA platform](https://imi-biomap.elixir-luxembourg.org/minerva/index.html?id=ISD_map_sidebyside_level&perfectMatch=false&modelId=359&backgroundId=588&x=2470.25&y=1545.75&z=4&overlaysId=1346)) that sustain the disease progression. Th cells sustain the progress of AD via expression of cytokines IL4, IL13, IL17A, IL22, IL31 and IFNG. While IL31 activates sensory nerve endings (SNEs) and promotes itch in AD, all other cytokines disturb KC proliferation and differentiation and stimulate release of more proinflammatory molecules (see “Th-derived interleukins effects on keratinocytes behavior and itch induction in AD” in the map [at the MINERVA platform](https://imi-biomap.elixir-luxembourg.org/minerva/index.html?id=ISD_map_sidebyside_level&perfectMatch=false&modelId=359&backgroundId=588&x=2470.25&y=1545.75&z=4&overlaysId=1346)). While disturbances in the KCs proliferation and differentiation cause the typical histopathology manifestations of AD, the KCs-derived proinflammatory molecules promote increased differentiation of T naïve cells into Th cells, sustaining skin inflammation. In AD, Th2 cells-derived interleukins also activate B cells and eosinophils (see “Th2 cells-activated B cells and eosinophils” in the map [at the MINERVA platform](https://imi-biomap.elixir-luxembourg.org/minerva/index.html?id=ISD_map_sidebyside_level&perfectMatch=false&modelId=359&backgroundId=588&x=2470.25&y=1545.75&z=4&overlaysId=1346)).

In PsO, initial trigger factors promote KCs to release proinflammatory molecules that activate plasmacytoid (pDCs) and conventional dendritic cell (cDCs). KCs also recruit neutrophils to epidermis in response to initial trigger factors. Some antigens can also be involved in the onset of PsO by directly activating pDCs (see “Keratinocyte-dependent or independent activation of dendritic cells and neutrophils in PsO” in the map [at the MINERVA platform](https://imi-biomap.elixir-luxembourg.org/minerva/index.html?id=ISD_map_sidebyside_level&perfectMatch=false&modelId=359&backgroundId=588&x=2470.25&y=1545.75&z=4&overlaysId=1346)). In response to these antigens and the KCs-derived proinflammatory molecules, pDCs express IFNA1 that, in turn, makes cDCs drive the differentiation of naïve T cells into specific Th cells and induce the activation of ILC3 cells (see “Dendritic cells-driven T cell polarization and ILC3 activation in PsO” in the map [at the MINERVA platform](https://imi-biomap.elixir-luxembourg.org/minerva/index.html?id=ISD_map_sidebyside_level&perfectMatch=false&modelId=359&backgroundId=588&x=2470.25&y=1545.75&z=4&overlaysId=1346)). Th cells sustain the progress of PsO via expression of IL17A, IL22 and IFNG. These cytokines disturb KC proliferation and differentiation and stimulate release of more proinflammatory molecules (see “Th-derived interleukins effects on keratinocytes behavior in PsO” in the map [at the MINERVA platform](https://imi-biomap.elixir-luxembourg.org/minerva/index.html?id=ISD_map_sidebyside_level&perfectMatch=false&modelId=359&backgroundId=588&x=2470.25&y=1545.75&z=4&overlaysId=1346)). Defects in KCs proliferation and differentiation cause the typical histopathology manifestations of PsO, and the KCs-derived proinflammatory molecules, such as TNF, promote increased differentiation of T naïve cells into Th cells, sustaining skin inflammation (see “TNF-mediated inflammatory circuit in PsO” in the map [at the MINERVA platform](https://imi-biomap.elixir-luxembourg.org/minerva/index.html?id=ISD_map_sidebyside_level&perfectMatch=false&modelId=359&backgroundId=588&x=2470.25&y=1545.75&z=4&overlaysId=1346)).

### Applications of the ISD map

In addition to be a graphical review of the molecular mechanisms of AD and PsO, we sought to verify if the ISD map could also be a hypothesis-generating resource via the discovery of potential mechanistic downstream effects of selected AD- or PsO-related genes of interest. For this purpose, we analysed the network structure of the map itself and the outcomes of the integration of the map with omics data (Figure 3).

**Figure 3.**
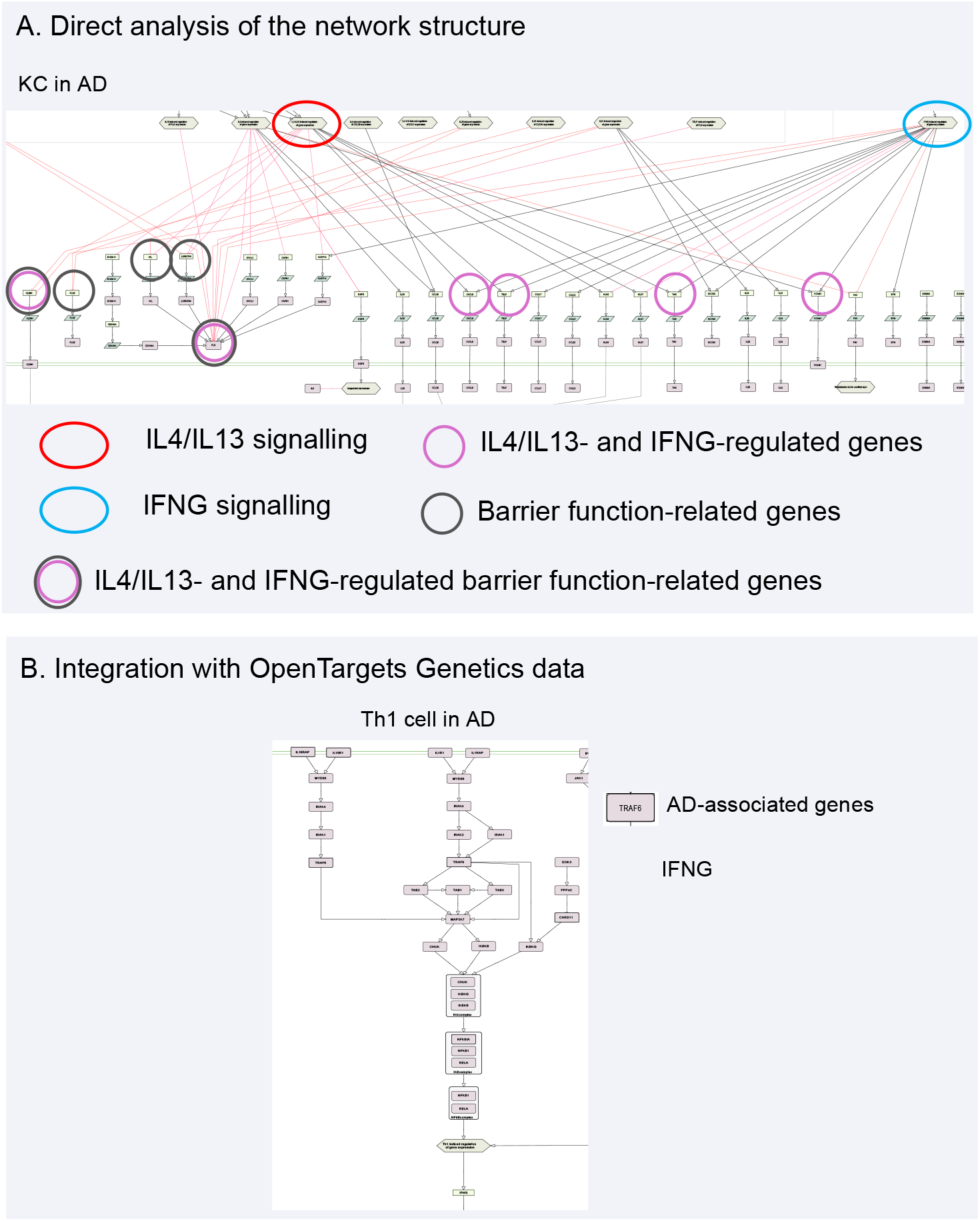
Possible applications of AD-derived omics data into ISD map: **A**. The direct analysis of network structure of the map allows users to identify that IFNG signalling is able to interfere with the IL4/IL13-regulated expression of several genes, including the ones involved in barrier function. **B**. By integrating AD-associated genes from the from Open Targets Genetics database into the map, it is possible to hypothesise that the regulation of IFNG expression in Th1 cells is heavily influenced by many upstream genes harbouring AD-associated SNPs.

### Direct analysis of the network structure

The direct analysis of the network structure per se may provide biological insights related to a disease of interest (15). To find compensatory pathways that could explain poor response of dupilumab, a widely used IL4R antagonist to treat moderate-to-severe AD (9), we analysed the network structure of the AD submap at both intercellular communication and intracellular (KCs and Th2 cells) levels (Figure 3A).

We could identify alternative pathways that explain, at least partially, the relatively low rate of remission following dupilumab treatment. In KCs, for instance, many genes involved in skin barrier homeostasis are downregulated not only by IL4/IL13 pathways, but also by IFNG, IL22, TSLP, IL-17A and IL25 signalling pathways (Figure 3A). So, the presence of these cytokines in skin could compensate for the inhibitory action of dupilumab on IL4R. It is noteworthy that an alternative pathway that potentially makes KCs partially insensitive to dupilumab is the one activated by IFNG as the expression of half of the IL4/IL13 pathway target genes are also modulated by the IFNG signalling pathway (Figure 3A).

### Omics data interpretation in AD and PsO maps

To show that the ISD map can be useful for supporting the discovery of potential AD- and PsO-related molecular mechanisms affected by genes or proteins that are prioritised via omics studies, we show here four use cases: (i) Integration of AD-associated genes from the Open Targets Genetics database (16) with the AD map, (ii) integration of PsO-associated genes from the Open Targets Genetics database with the PsO map, (iii) integration of differentially expressed proteins (DEPs) from protein expression profiles with the AD map and (iv) integration of differentially expressed genes (DEGs) from gene expression profiles with the PsO map.

#### Suggesting mechanistic consequences of gene variants

We collected genes harbouring variants (SNPs) associated with AD and PsO from the Open Targets Genetics database (16) (see Supplementary Methods in Supplementary Material for details). These AD- and PsO-associated genes were integrated with the ISD map as a publicly available dataset (for more details, refer to the user quick guide on how to explore and use the ISD map in the Supplementary Material). The AD map contains 28 of the 330 AD-associated genes, and the PsO map contains 42 of 794 PsO-associated genes.

We checked which inter- and intracellular activities in the AD and PsO maps were enriched in, respectively, AD- and PsO-associated genes. For this purpose, we performed a ISD map-specific pathway enrichment analysis (as described in the Supplementary Methods in the Supplementary Material): PsO-associated genes *versus* PsO map and AD-associated genes *versus* AD map. In AD map, intracellular activity of KCs (adjusted p-value = 0.0004) and intercellular activities of ILC2 cells (adjusted p-value = 0.0002), Th2 cells (adjusted p-value = 0.0030), mast cells (adjusted p-value = 0.0140) and basophils (adjusted p-value = 0.0326) were significantly enriched. Interestingly, the KC intracellular activity in the acute phase, but not in chronic disease stages, is significantly enriched (adjusted p = 0.0163) in AD-associated genes. This indicates that the KC-linked genetic component of AD is likely to influence the onset of AD and/or initial phases of disease activation.

Regarding PsO, KCs (adjusted p < 0.0001), Th17 cells (adjusted p = 0.0104) and γδ T cells (adjusted p = 0.0038) are significantly enriched in PsO-associated genes. As the main executor of the inflammatory circuit in psoriasis, KCs are expected to be enriched in PsO-associated genes. Th17 cells are also expected to be enriched as they play a pivotal role in keeping the IL23-IL17A/IL22 axis activated. The enrichment of PsO-associated genes in γδ T cells is surprising but concordant with the observed activation of the IL23-IL17A/IL22 axis in γδ T cells. Interestingly, while IL17A is not a PsO-associated gene in Open Targets Genetics database, many of its upstream regulators, such as IL12B, IL23A and IL23R, and proteins belonging to its downstream signalling, such as CARD14, NFKBIA, CHUK, among others, are encoded by PsO-associated genes.

We extended the pathway enrichment analysis performed after the integration of disease-associated genes to the ISD map to investigate their influence at the mechanistic level. To this end, we manually inspected the pathways of the ISD map for proteins encoded by the matched disease-associated genes that directly influence other proteins. As discussed previously, IFNG seems to partially compensate for IL4R inhibition by positively stimulating the expression of several AD-promoting genes also stimulated by IL4R in KCs (Figure 3A). As IFNG is mainly produced by Th1 cells, we checked the Th1 cell map for the presence of proteins encoded by AD-associated genes that could somehow influence IFNG expression. Interestingly, there are five proteins encoded by AD-associated genes (IL18RAP, IL18R1, TRAF6, CARD11 and NFKBIA) upstream to the IFNG expression (Figure 3B). We hypothesise that SNPs in these genes could favour IFNG expression in Th1 cells and, therefore, counteract the action of dupilumab, i.e., IL4R inhibition. Another example of mechanistic interpretation of the Open Targets Genetics data comes from the PsO map, specifically in KCs. In PsO, KCs are relatively resistant to cytokine-induced apoptosis (17). This resistance could be assigned, at least partially, to the presence of several proteins encoded by PsO-associated genes in apoptosis-regulating pathways. In fact, by exploring the map, we can see at least six proteins encoded by PsO-associated genes in these pathways: IFNG, INFGR2, TNFRSF1A, ESRRA, IRF1 and SOCS1. The most prominent pathway would be the one triggered by IFNG via IFNGR2 and IRF1 culminating in the expression of SOCS proteins. Remarkably, all proteins in this apoptosis-regulating pathway are encoded by PsO-associated genes and the underlying SNPs could favour the inhibition of apoptosis in KC (Figure 4A).

**Figure 4.**
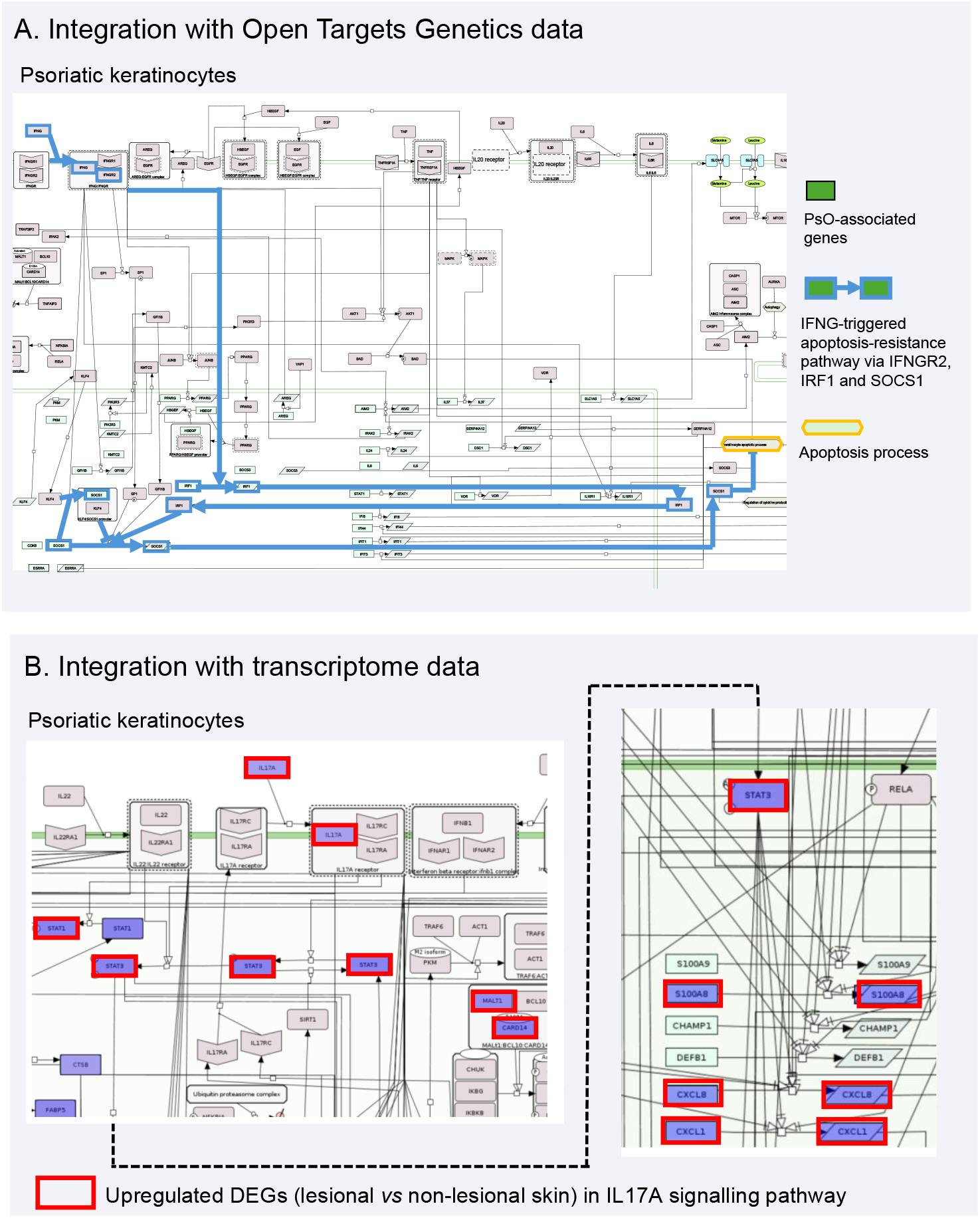
Possible applications of PsO-derived omics data into ISD map: **A**. By integrating PsO-associated genes from the from Open Targets Genetics database (dark green boxes) into the map, it is possible to hypothesise that the characteristic keratinocyte resistance to apoptosis observed in PsO is heavily influenced by many upstream genes harbouring PsO-associated SNPs. Blue thick lines represent causal interaction in the pathway from IFNG to the apoptotic process. **B**. The integration of differentially expressed genes (DEGs) from a meta-analysis derived (MAD) transcriptome of psoriasis comparing lesional and non-lesional skin samples reveals that not only IL17A is upregulated in PsO, but also signalling proteins and target genes downstream to IL17A in keratinocytes.

The above examples show that, through the integration of Open Target Genetics data with ISD map, we could determine the contextual relevance for AD- and PsO-associated genes and check if they fit into the existing molecular and cellular understanding of AD and PsO. Moreover, we were also able to formulate two hypotheses: resistance to dupilumab due to enhanced expression of IFNG in Th1 cells favoured by SNPs in upstream IFNG regulators and resistance to cytokine-induced apoptosis in psoriatic KCs due to altered upstream apoptosis regulators.

#### Suggesting mechanistic consequences of altered gene and protein expression profiles

We collected differentially expressed proteins (DEPs) from the study by He et al. (2020) (18) in which proteome expression profiles were measured in AD lesional and non-lesional stratum corneum samples taken from patients before and after treatment with dupilumab. From this study, we considered only DEPs calculated by comparing expression profiles of 353 inflammatory proteins extracted from lesional stratum corneum samples of patients before and after dupilumab exposure. Of the 132 dupilumab-induced differentially expressed inflammatory proteins (Dup-DEIPs), 20 could be found in the AD map. First, we performed an enrichment analysis to check if any intra- or intercellular activity was enriched in Dup-DEIPs in lesional stratum corneum. Only the intercellular activity of KCs (adjusted p-value = 0.0003) is significantly enriched in Dup-DEIPs. This is rather expected as stratum corneum is comprised virtually only by KCs. But, intriguingly, the intracellular activity of KCs is not significantly enriched; this can be explained by the fact that He and colleagues considered only a panel of inflammatory proteins for measuring expression; the intracellular pathways of KCs in the AD map contain 145 proteins and only 34 of them are classified as inflammatory.

Despite this limitation, we sought to check the KCs intracellular activity for finding which and how these Dup-DEIPs are distributed in KCs intracellular pathways. Of the 34 inflammatory proteins in KCs, nine (IL1RL1, IL17RA, IKBKB, CASP3, CASP8, CXCL8, CCL17, TNF and MMP9) are Dup-DEIPs, all being downregulated by dupilumab. We first checked which of these Dup-DEIPs are regulated by IL4/IL13 signalling. Of these nine proteins, only CXCL8 is a IL4/IL13 target and, as expected, it is downregulated. While CXCL8 is downregulated by dupilumab, the expression of TSLP, another IL4/IL13-induced inflammatory protein, seems not to be affected; TSLP is also regulated by IFNG signalling according to the map, so the TSLP expression could be rescued by IFNG in the absence of an active IL4R. IL17RA is downregulated by dupilumab and, therefore, we would also expect a downregulation of IL17RA signalling inflammatory target proteins. However, the DEP data provide no evidence that either of its inflammatory targets in KCs, namely CCL20, CSF3 and IL33, are affected by dupilumab. This suggests alternative pathways — such as IFNG for IL33, IL26 for CCL20 and an unknown pathway for CSF3 — that rescue the expression of these proteins. Finally, it is possible to realize that, via this integration with proteomic data, molecular connections between IL4R and the above-mentioned Dup-DEIPs — except for CXCL8 — are still missing in the AD map. This can be due to either AD knowledge not yet captured by biocurators or a real knowledge gap concerning such connections.

We collected DEGs from a meta-analysis derived (MAD) transcriptome of psoriasis comparing lesional and non-lesional skin samples published by Tian et al. in 2012 (19). From this study, we integrated into the map differentially expressed genes (DEGs) belonging to the set built by combining the results of five microarray data sets (MAD5) where the transcriptome profiles of lesional and non-lesional psoriatic skins were compared. Of 1116 DEGs, 63 mapped to proteins in the PsO map. By performing an enrichment analysis to check which intra- or intercellular activity are enriched in DEGs, we found that both inter-(adjusted p-value < 0.0001) and intracellular activities of KC (adjusted p-value < 0.0001) are enriched in DEGs, as well as intercellular activity of neutrophils (adjusted p-value = 0.0003). These findings are not surprising and serve the purpose of reinforcing the vital roles these cells play for psoriasis progression. The significant enrichment in melanocyte activity (adjusted p-value = 0.0410) and the lack of enrichment in Th17 cell activity are more surprising. While Abdel-Naser et al. (2016) showed that lesional psoriatic skin has an increased activity and number of epidermal melanocytes, the number of Th17 cells might not be different between non-lesional and lesional skin (20). This may at first appear inconsistent with the experimental evidence that IL17A is upregulated in lesional skin, but by inspecting the PsO map we can observe that IL17A is also produced by neutrophils, ILC3 cells, gamma-delta T cells, mast cells and Tc17 cells in addition to Th17 cells. As IL17A positively influences KC expression of mainly inflammatory proteins, we sought to verify whether IL17A downstream signalling and target proteins in KCs were also upregulated. Indeed, some of the IL17A downstream signalling proteins, e.g., STAT1 and STAT3, and target proteins, e.g., S100A8, CXCL1 and CXCL8, are upregulated. Interestingly, MALT1 and CARD14, signalling proteins that boost the NFKB signalling pathway — a pathway that is an important drive for IL17A signalling — are also upregulated (Figure 4B).

The above examples show that, through the integration of omics data with ISD map, we could (1) formulate hypotheses, i.e., IFNG rescues expression of dupilumab-downregulated proteins, and (2) identify gaps in the map or even in the AD-related knowledge itself that warrants further experimental investigation.

### Probabilistic Boolean simulation of IFNG modulation of IL4/IL13 pathway in AD

Static disease maps, such as the ISD map, can be converted into dynamic models, i.e., Boolean networks, to enable *in silico* simulations and predictions (21). Previous work has demonstrated that parts of the COVID-19 and Parkinson’s disease maps could be used to draw translational inferences from Boolean network simulations (15)(22). Here, by converting the IL4/IL13 signalling cascade in AD KCs into a Boolean model, we analysed the impact of IFNG on the effectiveness of dupilumab, focusing on sensory perception of itch and the activity of proteins TSLP, KLK5 and KLK7. The models demonstrate that, after approximately 100 iterations, the levels of TSLP, KLK5, KLK7, and sensory perception of itch stabilize, indicating a steady-state response in the network under the given conditions (Figure 5 and Supplementary Figures S1 and S2 in the Supplementary Material). This stabilization point can be used as a reference for understanding the system’s behaviour and response to treatments. By modelling the effects of IFNG and IL4 on key dysregulated molecules in AD, we observe that the presence of IFNG (Figure 5) is associated with less effective control of sensory perception of itch. This may be due to sustained TSLP levels (Figure S1 in Supplementary Material) and persistent inflammation, which counteract the benefits of dupilumab. Conversely, the absence of IFNG (Figure S2 in Supplementary Material) allows for a significant reduction in TSLP, leading to better control of itch and improved patient outcomes (Figure S2 in Supplementary Material). These findings may give insights on the importance of targeting multiple cytokine pathways to optimize therapeutic strategies for AD.

**Figure 5.**
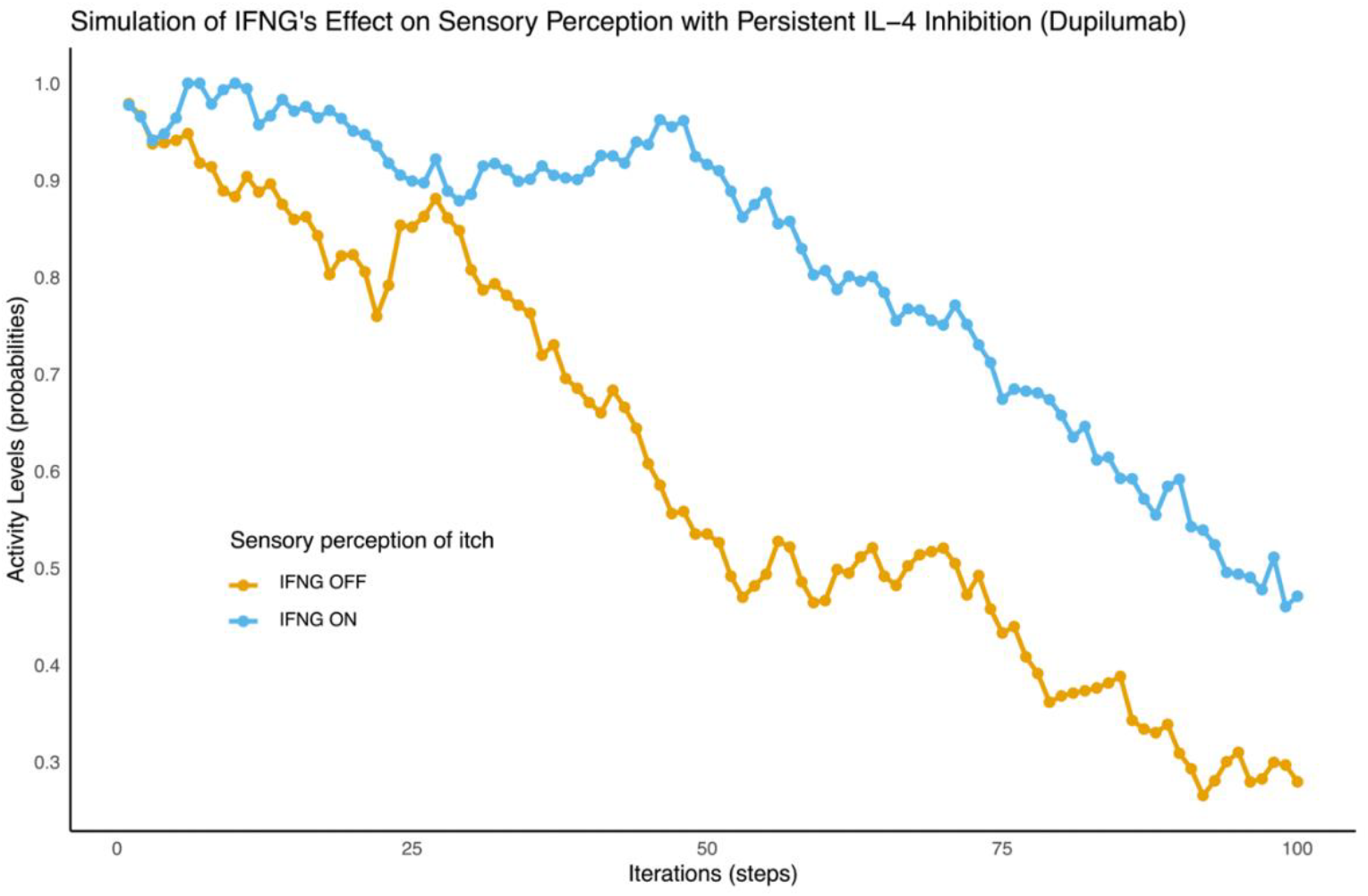
Boolean network simulation of IFNG effect on sensory itch perception after dupilumab treatment: Comparison between the decreasing sensory perception of itch (given by the activation probability as shown in ‘y’ axis) across time (in fact, pseudotime represented by iterations as shown in ‘x’ axis) when IFNG is expressed (“IFNG ON, IL4 OFF”) and not expressed (“IFNG OFF, IL4 OFF”) during persistent IL4R inhibition with dupilumab.

## DISCUSSION

The representation of biomolecular processes as generic networks, i.e., networks of unsigned directed or undirected interactions among biomolecules, has become popular in systems biomedicine (12). The analysis of these biomolecular networks can be useful, for example, for suggesting biomarkers and drug targets for disease. However, these networks are generic, implying that interactions among their elements do not necessarily reflect true interactions under the context of a specific disease or in a specific tissue.

By building a disease map, we move from a generic network characterization of a disease to a more disease-specific molecular mechanistic network, considering different cell types, tissues, organs and disease states (23). As a disease map is tailored to a specific disease, we assume that it can provide more mechanistic insights than the generic network. This assumption is supported by the fact that an increasing number of studies have reported methods to enhance generic networks specifically for a disease to prioritize drug targets and candidate biomarkers (24)(25)(26).

Based on the above-mentioned considerations and the previous successful applications of disease maps (27)(28)(29), we opted for the construction of the network representing the AD and PsO-specific molecular mechanisms in a disease map format, i.e., the ISD map. We could demonstrate that this map can be useful as an interactive, graphical tool for reviewing AD and PsO molecular mechanisms and identifying potential effects on altered pathways as well as generating mechanistic hypotheses.

### Strengths and limitations of the study

This map was constructed via manual curation of biomedical literature in a systematic search supplemented by domain expert recommendations, a laborious and time-consuming process that has resulted in deep coverage of key mechanisms of both AD and PsO. However, the molecular mechanisms present in the map cannot include all knowledge of AD and PsO-specific mechanisms thus reflecting the map coverage limited by the biocuration process. Data in the map also represent a snapshot in time, since the field of biomedical and molecular research is rapidly expanding. To further develop the map and keep it as updated as possible, periodic revisions are planned and the long-term sustainability of the ISD map will be guaranteed by the ELIXIR Luxembourg’s “Disease Map” service (https://elixir-luxembourg.org/services/catalog/minerva/).

## Notes

**Conflict of Interest: CHS:** received research funding from consortia with multiple industry partners (BIOMAP-IMI.eu, HIPPOCRATES-IMI.eu, PSORT.org.uk); Sanger/OPENTARGETS; Astrazenca; Boerhinger-Ingleheim. **JR, MN:** work for UCB and hold shares in UCB. The rest of the authors declare no competing financial interests.

### Competing Interest Statement

CHS: received research funding from consortia with multiple industry partners (BIOMAP-IMI.eu, HIPPOCRATES-IMI.eu, PSORT.org.uk); Sanger/OPENTARGETS; Astrazenca; Boerhinger-Ingleheim. JR, MN: work for UCB and hold shares in UCB. The rest of the authors declare no competing financial interests.

