## Supplementary Material for "The inflammatory skin disease map (ISD map): an interactive computational resource focused on psoriasis and atopic dermatitis molecular mechanisms"

### HOW TO NAVIGATE AND USE THE ISD MAP: QUICK USER GUIDE

#### Accessing the maps

The main entry point to the ISD map is available at https://imi-biomap.elixir-luxembourg.org/. This is a side-by-side simplified comparison of the contents of atopic dermatitis (AD) and psoriasis (PsO) maps.
To access the maps, follow the steps below.

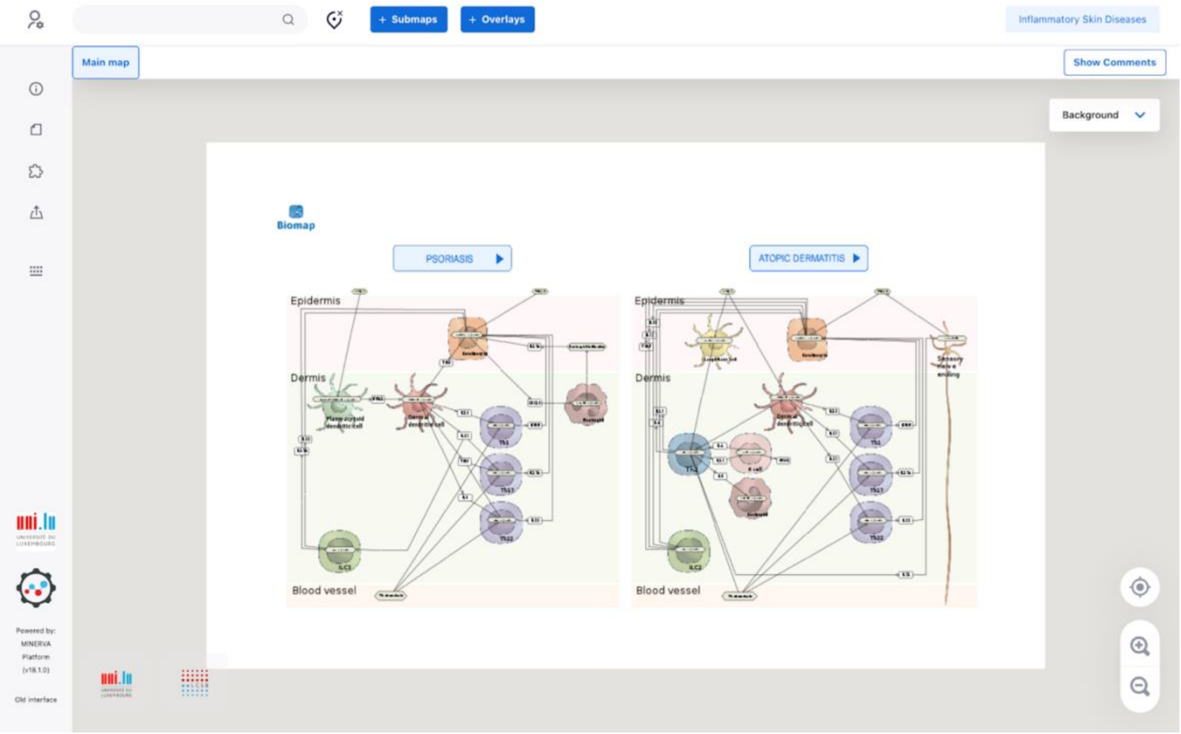


To access the individual, i.e., AD or PsO, intercellular communication maps, please click on the button with disease name above each part of the diagram.

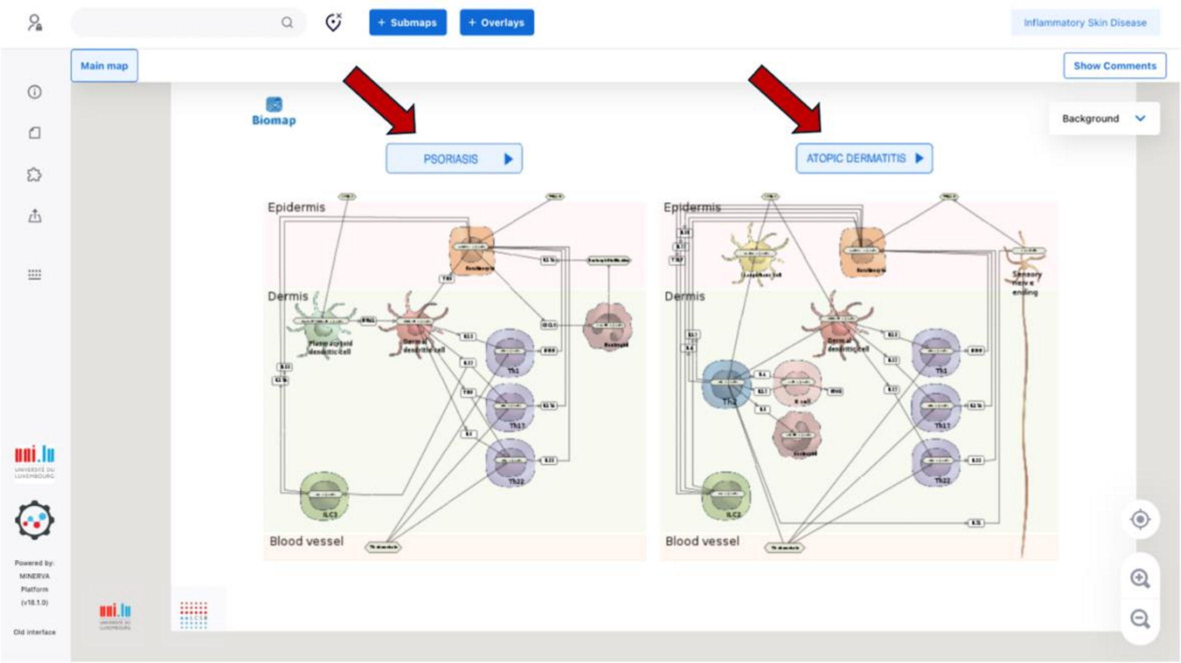


Clicking on the “ATOPIC DERMATITIS” button, you will be directed to the AD map.

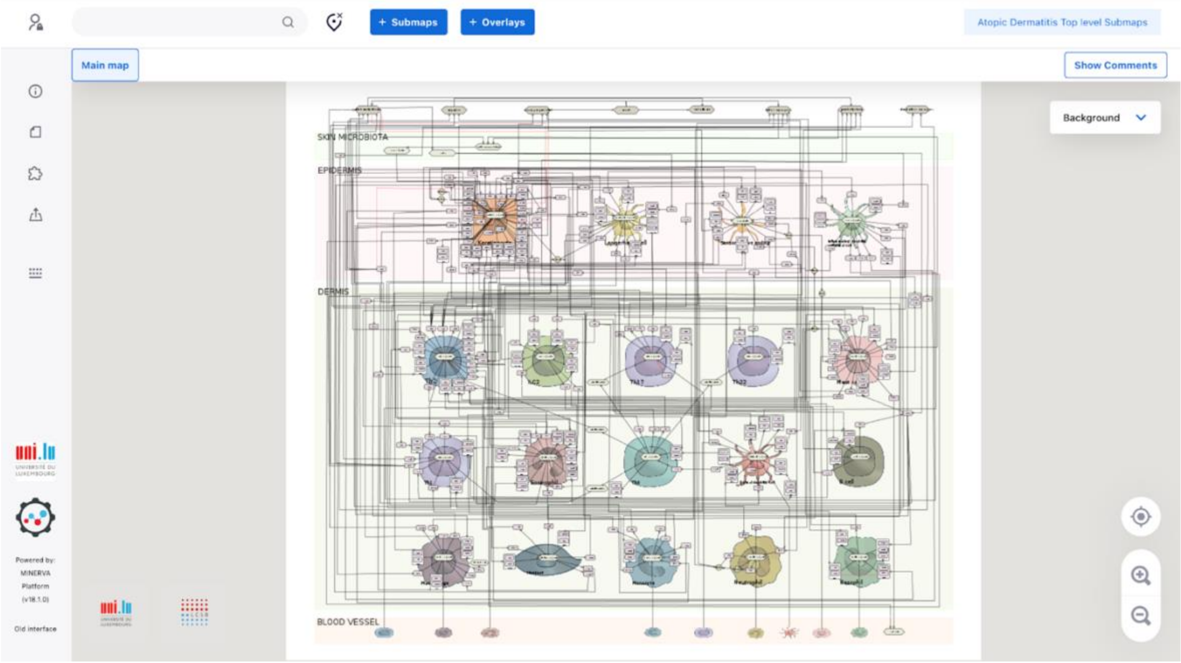


#### Exploring the maps

You can explore the map by navigating it similarly as you do in Google Maps. Use the buttons on the lower right corner to zoom in, zoom out and center the map content.

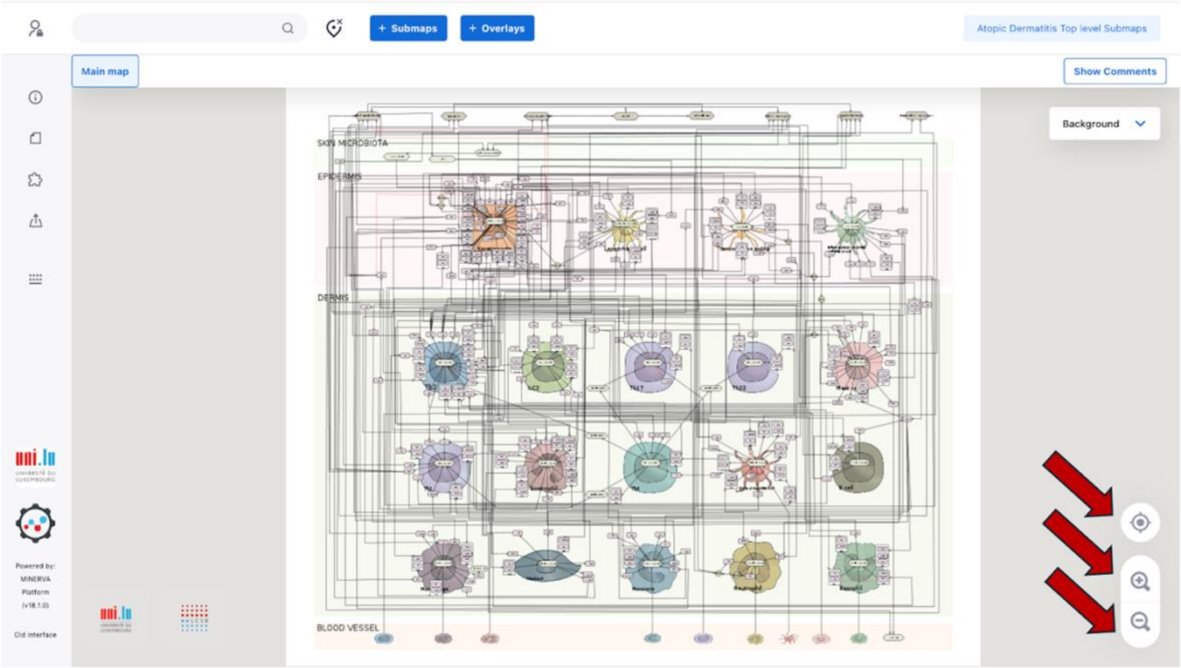


To access information about map elements - cell types, phenotypes, proteins, genes, interactions - click on the element, which becomes marked with a blue anchor, and check the left panel, which contains annotations referring to the element.

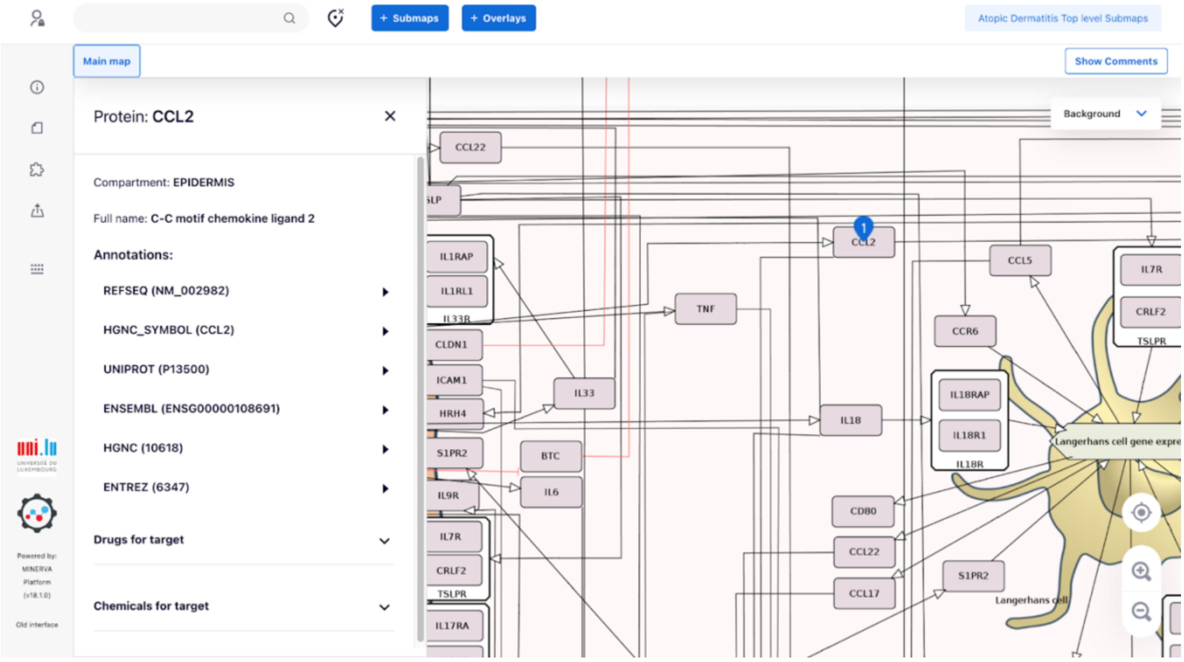


An arrow connecting elements on the map (a reaction) represents an interaction between them. By clicking on it, it becomes blue and the left panel displays references to publications supporting this reaction, and elements linked by the interaction.

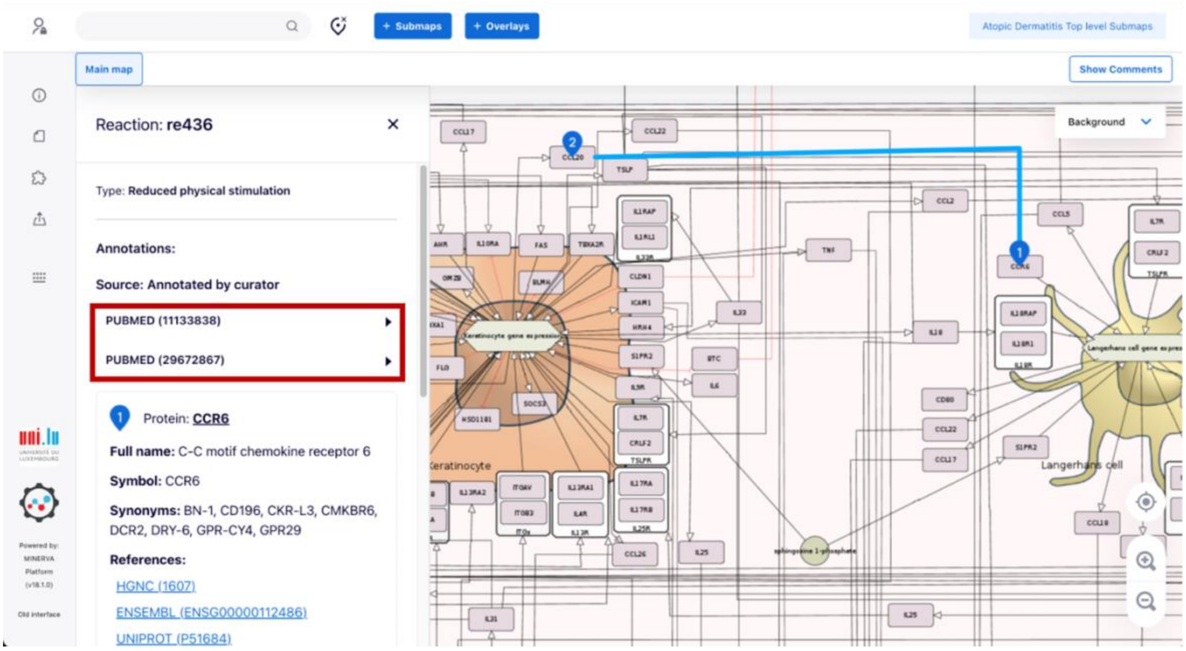


#### Search

You can search for elements using the search panel (as shown below). After clicking on 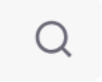 or hitting "Enter", the searched element will be marked with blue anchors in the map. Select "Perfect match" to focus search results.

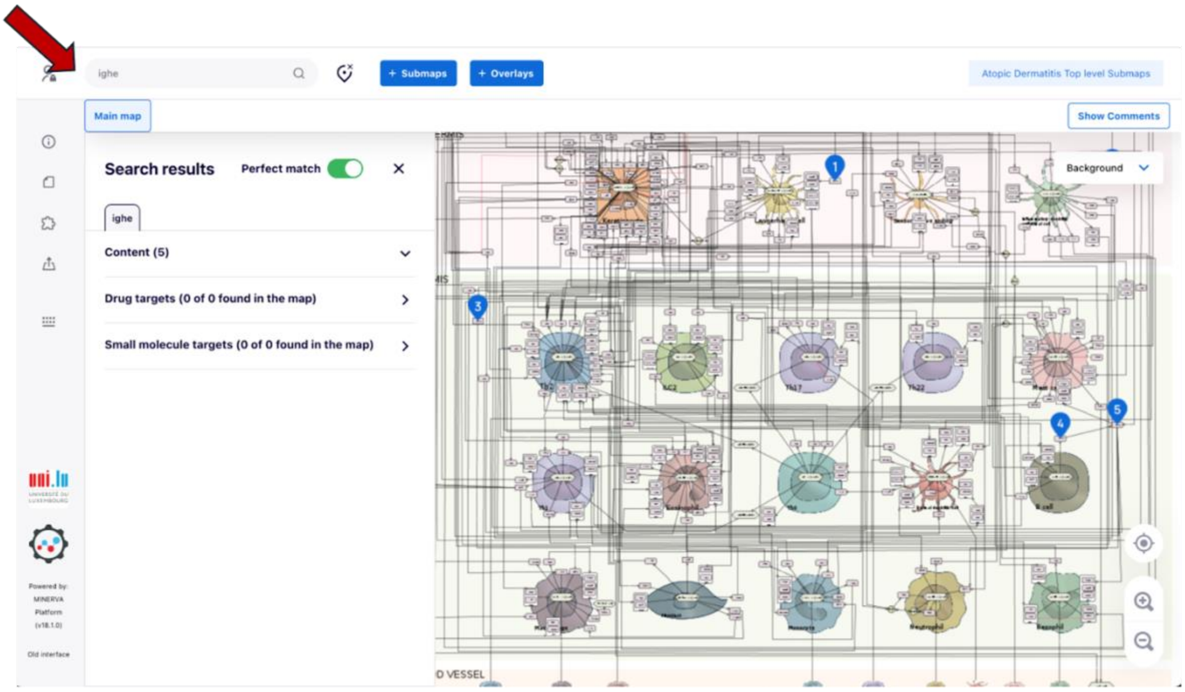

You can access submaps containing intracellular pathways in key cell types in the intercellular communication map. This can be done by clicking 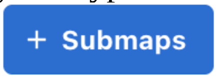

, then selecting the submap of interest among the listed ones in the panel appearing in the left side, and finally clicking “ >” in the same row of the selected submap.

[Figure is coming soon]

For example, by clicking “>” in the same row of the submap “Keratinocyte”, you will be directed to the submap containing AD-related intracellular pathways in keratinocytes. If you would like to have a broader view of the submap, close the “Submaps” panel by clicking on “X”.

[Figure is coming soon]

**IMPORTANT:** you can perform exploration and search activities in submaps.

#### Data visualisation

Click 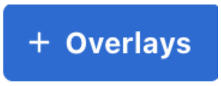

to visualise i) genes harboring AD- or PsO-specific SNPs , ii) differentially expressed genes and proteins with their normalized log fold change values and iii) AD- and PsO-linked biological processes discussed in the paper. The left panel will display the abovementioned list . Click “View” to visualise an overlay of interest, e.g., *Omics: Differentially expressed proteins from dupilumab vs untreated lesional skin He et al (2020)*.

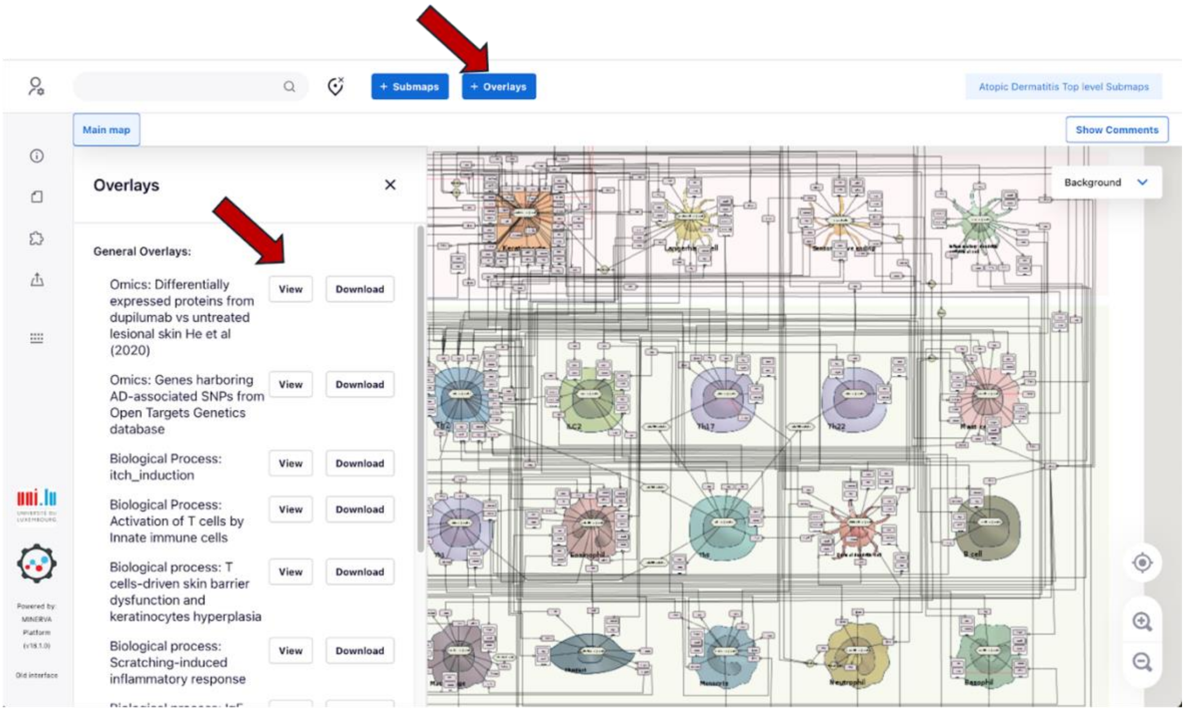


After clicking "View", the elements present in the selected overlay, which in this case contains quantitative omics data, are highlighted in a gradient color according to their normalized log fold change as shown in the horizontal bar at the bottom of "Overalys". If you want to remove the highlights, click "Hide".

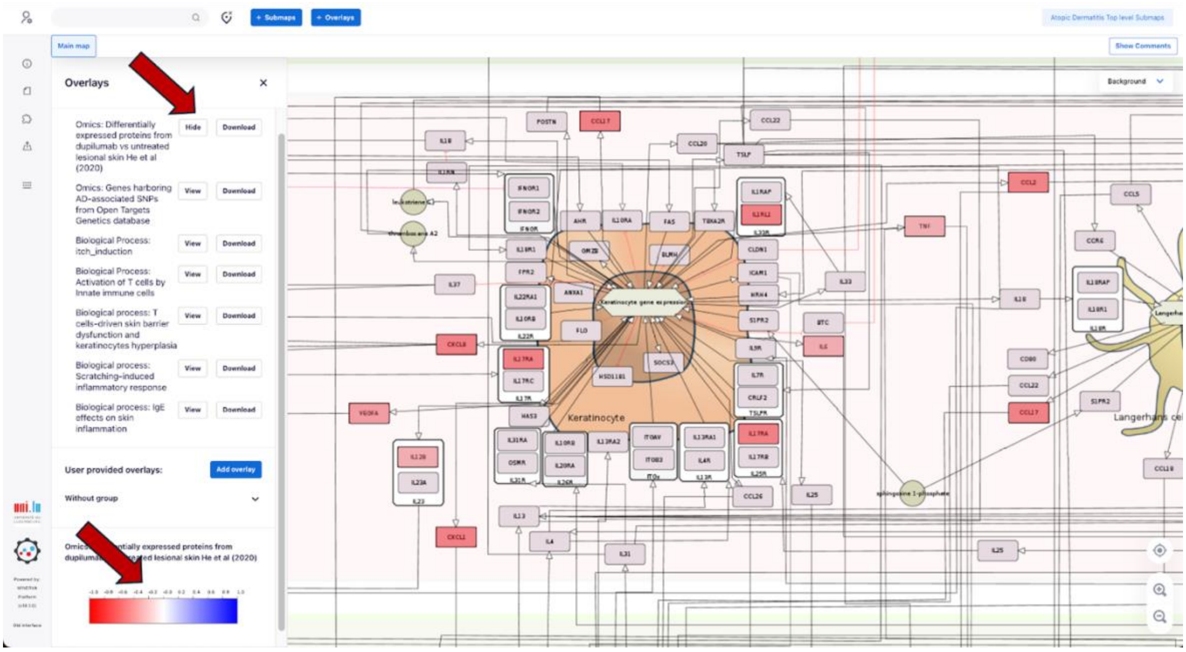


If you selected an overlay containing qualitative omis data "View", the elements present in the selected overlay, which in this case contains quantitative omics data, are highlighted in a gradient color according to their normalized log fold change as shown in the horizontal bar at the bottom of “Overalys”. If you want to remove the highlights, click “Hide”.

### SUPPLEMENTARY METHODS

#### Map construction and availability

Causal interactions relevant to AD and PsO were extracted from biomedical literature and represented in a diagrammatic visualisation, referred to as the map, using the CellDesigner tool (1) and following the Systems Biology Graphical Notation (SBGN) standard (2) (https://sbgn.github.io/learning). The only exception was in the representation of drugs, where we utilized the CellDesigner-specific glyph. The ISD map was subsequently uploaded to the MINERVA platform (3) (https://imi-biomap.elixir-luxembourg.org/minerva/) for straightforward access and exploration. Further details are provided below.

#### Capturing AD- and PsO-related causal interactions from literature

The ISD map was developed by adhering to many recommendations from the recently published guidelines for constructing disease maps (4). Initially, we manually curated reviews and original peer-reviewed research articles suggested by domain experts from the BIOMAP consortium. While most of these expert-selected papers focused on experimentally validated causal interactions specific to AD and PsO, some described experimentally validated immune processes in general, which were included to fill gaps in disease-specific information. Most causal interactions were assigned to compartments (such as organs, tissues, cells, and organelles) based on information provided in the papers; in certain cases, this assignment was inferred using common knowledge about the usual locations where these interactions occur. We also documented details related to causal interactions (e.g., gene variants, phosphorylated or citrullinated residues in proteins) when available in the literature.

#### Encoding causal interactions into diagrams

The identified causal interactions were encoded into a diagrammatic visualization format using the CellDesigner tool (1), following its standard visual syntax, the CellDesigner Systems Biology Markup Language (SBML) extension. This encoding process adhered to the Systems Biology Graphical Notation (SBGN) standard (2) (https://sbgn.github.io/learning). We utilized both the SBGN Process Description (PD) and Activity Flow (AF) languages. In PD, a causal interaction is depicted as a state transition of a biochemical entity (the regulated entity), including biochemical details about the type of molecular transition and how it is influenced by a regulator entity. In AF, a causal interaction is shown as a simple directed link between two entities, indicating the type of regulation, such as activation or inhibition

#### Identification and annotation of the map elements

To ensure compatibility with systems biology tools and external databases, all entities (proteins, RNAs, genes, complexes, metabolites, drugs, and phenotypes), compartments (organelles, cells, tissues, and organs), and interactions in the map were thoroughly annotated. This process followed the Minimal Information Requested in the Annotation of Models (MIRIAM) guidelines (5), a standard for annotating and curating computational models and maps. Proteins, RNAs, and genes were identified using their official symbols from the HUGO Gene Nomenclature Committee (HGNC, https://www.genenames.org), allowing the MINERVA platform, which facilitates visualization and navigation of the map (see details below), to automatically add further annotations such as Ensembl, Entrez Gene, RefSeq, and UniProt IDs. If HGNC could not recognize certain symbols or terms, additional annotations were manually assigned. Complexes were annotated, when available, to Gene Ontology (GO) cellular component terms and their corresponding IDs (6) (7), while metabolites were identified by their ChEBI names (8) (https://www.ebi.ac.uk/chebi) and IDs. Phenotypes were identified using GO biological process terms and their IDs (http://geneontology.org/docs/ontology-documentation) (6) (7) for biological processes, or Medical Subject Headings (MeSH) terms and their IDs (https://www.ncbi.nlm.nih.gov/mesh) for disease-related elements. Annotations not automatically retrieved by MINERVA were added manually via the MIRIAM section of CellDesigner. References for interactions, including PubMed IDs of the source papers, were annotated using the relation "bqbiol: isDescribedby", while other annotations were added using the "bqmodel:isEncodeby".

##### Map availability and visualisation in the MINERVA platform

The ISD map is accessible as an online interactive map via the Molecular Interaction Networks Visualization (MINERVA) platform (3) (https://imi-biomap.elixir-luxembourg.org/minerva/). MINERVA is a standalone web server designed for the visual exploration, analysis, and management of molecular networks encoded in systems biology formats such as CellDesigner, SBML, and SBGN. It offers automated content annotation and verification, along with features like overlaying experimental data (e.g., transcriptomics, gene variant data) on the visualized networks. For more information on MINERVA's functionalities, please refer to the documentation at https://minerva.pages.uni.lu/doc/.

#### Integration, visualisation and analysis of omics data

##### Integration of Open Targets Genetics data to the map

For the analysis of possible mechanistic effects of gene variants on downstream molecular processes in ISD map, we collect AD- and PsO-associated genes, i.e., genes harbouring variants, such as single nucleotide polymorphisms (SNPs), associated with AD and PsO, respectively, from the Open Targets Genetics database (9). We collected these genes on 26.06.2024 by using the Experimental Factor Ontology (EFO) identifiers of AD (EFO_0000274) and PsO (EFO_0000676). We prepared MINERVA-compatible files for the creation of visual overlays of these variants in ISD map in the MINERVA platform: the "AD OT Genetics" overlay for the AD map (e.g., https://imi-biomap.elixir-luxembourg.org/minerva/index.xhtml?id=AtoDTopSubmaps1-07-24, "General Overlays" tab) and the "PsO OT Genetics" overlay for the PsO map (https://imi-biomap.elixir-luxembourg.org/minerva/index.xhtml?id=PsO_02072024, "General Overlays tab").

##### Integration of proteomics and transcriptomics data to the map

For the analysis of omics data integration with the map, we used two datasets: one proteomic dataset related to AD (10) and a transcriptomic dataset related to PsO (11).

The AD-related proteomic data was collected from the study by He et al (10); in this study, inflammatory proteome expression profiles (based on a panel of pre-selected 353 inflammatory proteins) were measured via Olink proteomic assay in AD lesional and non-lesional skin samples taken from patients before and after treatment with dupilumab. For our integration, we considered only differentially expressed proteins (DEPs) calculated by comparing expression profiles extracted from lesional skin samples of patients before and after dupilumab exposure. DEPs with FDR < 0.05 and fold-change (FCH) > 1.5 were collected from the paper's Supplementary Table E2, specifically from columns A (HGNC symbol of the proteins), H ("FCH LS Post-Rx versus Pre-Rx") and J ("FDR LS Post-Rx versus Pre-Rx"). We then normalized the FCH of DEPs to the [-1,1] range. Finally, we prepared the file for the creation of a visual overlay of these DEPs in ISD map in the MINERVA platform (https://imi-biomap.elixir-luxembourg.org/minerva/index.xhtml?id=AtoDTopSubmaps1-07-24), "General overlays" tab, ", "AD Inflammatory Proteome Dupilumab").

The PsO-related transcriptomic data was collected from the study by Tian et al (11); in this study, differentially expressed genes (DEGs) were defined via a meta-analysis of five transcriptomic studies (GSE6710, GSE11903, GSE14905, GSE13355 and (12) comparing lesional and non-lesional skin samples (the MAD-5 transcriptome). For our integration, we considered only DEGs with FDR < 0.05 and fold-change (FCH) > 2 extracted from the Supplementary Table S2. We then normalized the FCH of DEGs to the [-1,1] range and prepared the file for the creation of a visual overlay of these DEGs in ISD map in the MINERVA platform (https://imi-biomap.elixir-luxembourg.org/minerva/index.xhtml?id=PsO_02072024), "General overlays" tab, "PsO MAD5").

#### Functional enrichment analysis in MINERVA

To check if some of intra- or intercellular activities in both ISD map were enriched in AD- and PsO-related omics-prioritised genes, we used the MINERVA's GSEA plugin (13) as described in details here: https://minerva.pages.uni.lu/doc/plugins/gsea-plugin/. In brief, the MINERVA's GSEA plugin considers as the background gene list all genes present in the map; as the pathway database source, the plugin considers the list of annotated pathways in the own map. The statistical test used is the hypergeometric test and the obtained p-values are adjusted for multiple comparisons by using the Bonferroni test. Enriched pathways were those with adjusted p-value < 0.05.

#### Boolean network and simulation

We used Probabilistic Boolean Modelling (PBM) to simulate the effects of IFNG on sensory perception of itching after treatment with dupilumab, i.e., inhibition of IL4R. For this purpose, we first converted the individual KC map in the AD map into a Boolean network (BN) in an automated fashion using CaSQ tool (14). Then we considered the following pathways in the converted BN for our simulations: i) IL4R -> TSLP -> sensory perception of itching, (ii) IL4R -> KLK5 -> sensory perception of itching, (iii) IL4R -> KLK7 -> sensory perception of itching, (iv) IFNG -> TSLP -> sensory perception of itching, (v) IFNG -| KLK5 -> sensory perception of itching and (vi) IFNG -| KLK7 -> sensory perception of itching. The PBM approach uses a series of random walks to determine the probability of components within the model (15). This approach integrates qualities of both discrete and continuous Markov processes within a Monte Carlo framework (16). To establish a foundational baseline for our simulations, we parameterized initial state probabilities of ON/OFF. This step is critical as it sets the starting point for the model, reflecting the pre-simulation status of molecular interactions. The equation for updating the state probabilities is given by:

$$Pt + 1(s) = \sum s' \in SPt(s') \cdot P(s'|s)$$

where Pt+1(s) is the probability of state s at the next time point and P(s′|s) is the transition probability from a previous state s′ to the current state s.

### SUPPLEMENTARY RESULTS

#### The ISD map contents: a graphical review of mechanisms

As mentioned in the main text, the ISD map was built based on the review of AD- and PsO-related articles and, as such, the exploration of the contents of this map leads naturally to a graphical review of established molecular mechanisms of these diseases. While in the main text we summarise the main molecular and cellular aspects of AD and PsO, here we summarise the mechanisms at the intercellular and intracellular levels. As in the main text, we also provide here hyperlinks that allow readers directly access some of the mentioned region of the ISD map.

#### Intercellular communication in AD and PsO

The intercellular communication view for AD includes 19 cell types interconnected by cytokines and their receptors (see the map at the MINERVA platform). These are KCs, LCs, SNEs and inflammatory dendritic epidermal cells (IDECs) in the epidermis, and Th and B cells, granulocytes, cDCs, monocytes, macrophages, ILC2 cells and fibroblasts in dermis. The activation of this network in AD starts with the disruption of the skin barrier homeostasis, leading to a release of alarmins and allowing foreign substances to encounter LCs, cDCs and IDECs. In response, these cells release interleukins and chemokines that promote differentiation and recruitment of Th2, Th1, Th17 and Th22 cells (17) (see "Activation of T cells by innate immune cells" in the map at the MINERVA platform). Th cell interactions drive (i) the emergence of epidermal hyperplasia, lichenification, sensory perception of itch, scratching and inflammatory response and (ii) the maintenance of the skin barrier dysfunction.

Lichenification, one of the AD hallmarks, is characterised by epidermal hyperplasia, i.e. pathological multiplication of KCs. It emerges from stimulatory effects of KCs by IFNG, IL17A and IL22 released, respectively, by Th1, Th17 and Th22 cells. IL17A also plays a role in the maintenance of skin barrier dysfunction via downregulation of proteins BTC, CLDN1 and FLG in KCs. Th2 cells-derived granzyme B (GZMB) also takes part in skin barrier dysfunction (see "T cells-driven skin barrier dysfunction and keratinocytes hyperplasia" in the map at the MINERVA platform). Both skin barrier dysfunction and lichenification are influenced by scratching, promoted by an increased sensory perception of itch. This increase perception of itch, in turn, is generated by an overstimulation of sensory nerve endings (nociceptor sensory neurons) in response to a combination of IL4, IL13 and IL31 derived from Th2 cells, KC-derived TSLP, histamine released by mast cells, eosinophils-derived IL5-induced BDNF – a neurotrophin that promotes sensory nerve branching - and the expression of serotonin receptor HTR7 in SNEs and histamine receptors HHR1 and HHR4 in Th2 cells (see "Itch induction" in the map at the MINERVA platform). Itching elicits scratching, a behaviour that stimulates an AD-associated inflammatory response via IL4, IL13 and IL5 released by Th2 cells,

IL17A released by Th17 cells and mast cell-derived GZMB. These proteins influence the activities of KCs, B cells, fibroblasts, eosinophils, Langerhans cells, and SNEs (see "Scratching-induced inflammatory response" in the map at the MINERVA platform). Importantly, the elevated production of IgE (IGHE), a hallmark in AD, is due to IL4 and IL13 stimulation of B cells. IgE induces the production of interleukins and chemokines in mast cells and may trigger LCs, cDCs, macrophages, basophils and eosinophils to produce interleukins and chemokines, contributing to the continued skin inflammation (see "IgE effects on skin inflammation" in the map at the MINERVA platform).

The intercellular communication view for PsO is comprised by 17 different cell types interconnected mainly by cytokines and their receptors that are currently known to be involved in psoriasis: KCs, LCs, cDCs, plasmacytoid dendritic cells (pDCs), T cytotoxic (Tc), Th and B cells, granulocytes, ILC3 cells, melanocytes and fibroblasts (see the map at the MINERVA platform). The activation of this PsO-related cellular network can be linked to skin injury, causing KCs to release double-stranded DNA (dsDNA) and RNA (dsRNA) fragments complexed with antimicrobial peptides, notably CAMP (LL37). These complexes can stimulate several cells, including KCs themselves, neutrophils and antigen-presenting cells (APCs) such as LCs, cDCs and pDCs. These stimulated cells express, mainly via TLR signalling pathways, proinflammatory molecules able to recruit and activate both innate and adaptive immune cells. Neutrophils respond to the dsRNA-CAMP complex by secreting neutrophil extracellular traps (NETs) with IL17A, a cytokine playing a critical role in the pathogenesis of psoriasis (18). In addition to IL17A, NETs also contain other proteins, that promote inflammatory responses in KCs; these cells, in turn, produce neutrophil-attracting chemokines, such as CXCL8, CXCL1 and LCN2 (20) (21), which leads to more neutrophils infiltrating the epidermis. In addition, as NETs also contain dsRNA and CAMP, neutrophils are further activated and more NETs are released (22), creating an early self-amplifying inflammation cycle (see "Early innate immune-led self-amplifying inflammation cycle" in the map at the MINERVA platform).

In addition to this early KCs-neutrophils crosstalk, the activation of pDCs is essential for the development of PsO (23). Once activated, pDCs produce IFNA1 that stimulates cDCs to express TNF, IL1B, IL6 and IL23 (24). While TNF acts on cDCs, LCs and KCs by promoting the expression of proinflammatory proteins, including CCL20, the chemoattractant of the Tc17 and Th17 cells (25), cDCs-derived IL1B, IL6 and IL23 act together to stimulate differentiation of Tc17 and Th17 cells with subsequent expression of IL17A, IL17F and IL22 (26). These interleukins drive KCs to express more proinflammatory proteins, including chemokines, such as TNF and CCL20, and promote KCs hyperproliferation. This sustained cDCs-Tc17-Th17-KCs crosstalk creates an additional self-amplifying inflammation cycle in PsO (see "Innate to adaptive immune self-amplifying inflammation cycle" in the map at the MINERVA platform). This cycle can also include LCs, γδ T, ILC3 and mast cells, as they produce IL17A and IL22 and, except for LCs, are also attracted to skin via KC-derived chemokines (26). Finally, besides KCs, there are two other non-specialised immune cells: melanocytes and fibroblasts. Melanocytes are targets of not only TNF, but also Tc17 cells in PsO. Melanocytes express major histocompatibility (MHC) class I proteins that present ADAMTSL5 fragments in their surface, targeted for a noncytotoxic Tc cell–mediated autoimmune response (27). In turn, fibroblasts play a dual role in PsO. First, by expressing chemerin (RARRES2), they attract pDCs to skin (28) in the earliest phases of the disease. Second, by expressing tenascin C (TNC), they promote neurite outgrowth (29).

#### Intracellular mechanisms of AD and PsO

***Role of non-specialised immune cells in AD and PsO***

In AD and PsO, KCs are the most important non-specialised immune cells. KCs are involved not only in the onset of AD and PsO, but also in the maintenance of these two ISDs (30) (30). Although they are non-bone marrow-derived cells, KCs play a critical role in innate immune reactions and participate in the activation of adaptive immunity in the skin (32). Both AD and PsO maps show this aspect: KCs respond to a variety of endogenous and exogenous ligands and multiple interleukins derived from innate and adaptive immune cells by expressing chemokines and interleukins that attract and activate both innate and adaptive immune cells. In both diseases, this response takes place through the JAK-STAT, MEK-ERK and NFKB signalling pathways as we can observe in both AD and PsO maps. In AD, the chemokines CCL17 and CCL22, expressed in response to IFNG via JAK-STAT, MEK-ERK and NFKB pathways attract Th2 cells to dermis, while interleukin TSLP, expressed in response to TLRs, IL4/IL13 and IFNG, activates Th2 cells. In this case, KCs bridge two different adaptive immune cells, i.e., Th1 and Th2 cells. In PsO, KCs bridge innate and adaptive immune cells (e.g., ILC3 cells and Th17 cells) and an innate immune cell (neutrophil). For instance, CXCL8, a neutrophil chemoattractant, is produced in response to IL17A via the NFKB signalling pathway.

***T cell activity in AD and PsO***

At least five types of T cells - γδ T, Th1, Th2, Th17 and Th22 cells - play a role in AD or PsO. Here we describe in detail the intracellular pathways in Th1 and Th2 cells in AD and Th17 and γδ T cells in PsO.

Th2 cells are key in AD and are involved in both acute and chronic phases of this disease (33). As can be observed in the AD map, in response to IL33, IL25, TSLP and IL4 stimulation, Th2 cells express interleukins known to influence skin barrier dysfunction (IL4, IL13), eosinophil activation (IL5), keratinocyte differentiation and sensory perception of itch (IL4, IL13, IL31). IL33 promotes the activation of four transcription factors - ATF2, GATA3, JUN and the NFkB complex - via the MYD88-TRAF6-TAK1(MAP3K7)-TABs (MTTT) signalling cascade. Downstream p38 MAPK (MAPK14) activates ATF2 and GATA3. In parallel, downstream JNK signalling cascade activates JUN. Finally, parallel downstream IKKs activate the NFkB complex. IL25 also promotes the activation of the NFkB complex via TRAF6-TAK1(MAP3K7), but with TRAF3IP2 (ACT1) as the adaptor protein. Additionally, IL25 also activates a JAK-STAT signalling pathway to regulate downstream genes, as well as TSLP and IL4.

As for Th2 cells, Th1 cells also play a role in both phases of AD (32). In response to KCs-derived IL1B, Th1 cells express mainly IFNG, a proinflammatory protein that negatively influences KCs differentiation and positively influences inflammatory response (34). IFNG is expressed because of the activation of the NFkB complex via the MTTT cascade - the same used by IL33 in Th2 cells as previously discussed. IFNG is also upregulated by an autocrine loop in which IFNG itself stimulates IFNG expression via JAK-STAT signalling and the transcription factor TBX21.

Like Th2 cells in AD, Th17 cells in PsO are considered one of the most critical pathogenic factors (34). As a response to IL1B, IL6 and IL23 stimulation, Th17 cells express, among other proteins, IL17A and IL22 that, in turn, stimulate KC proliferation and expression of inflammatory proteins (36). IL1B binds to IL1RI and induces expression of the transcription factor IRF4 that, in turn, upregulates expression of RORC. The RORC-dependent transcription is then activated by IL6 and

IL23 via their receptors and JAK-STAT3 signalling. RORC in turn can induce transcription of IL17A, IL17F, IL21, IL22, IL23R, as well as CCR6, a chemokine that influences Th17 cell migration to sites of inflammation (37). STAT3 itself is also able to directly induce expression of IL17A, IL17F and IL22.

### SUPPLEMENTARY FIGURES

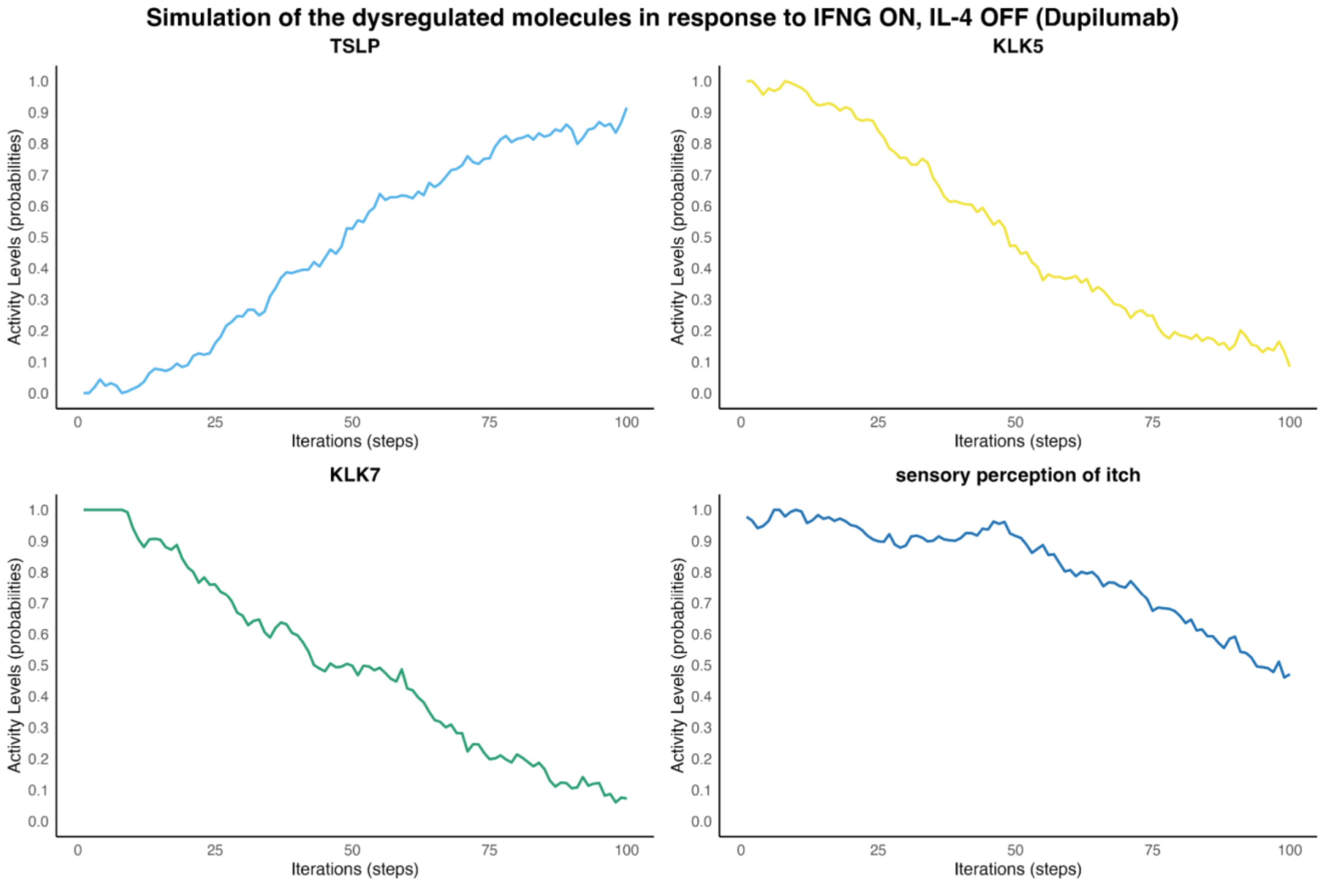


Figure E1. **Boolean network simulation of IFNG effect on outcomes after dupilumab treatment:** When IFNG is expressed (IFNG ON) in the presence of dupilumab (IL4 OFF), we observe an increase in TSLP levels, a decrease in KLK5 and KLK7 levels and a reduction of around 50% in the activity level of sensory perception of itch.

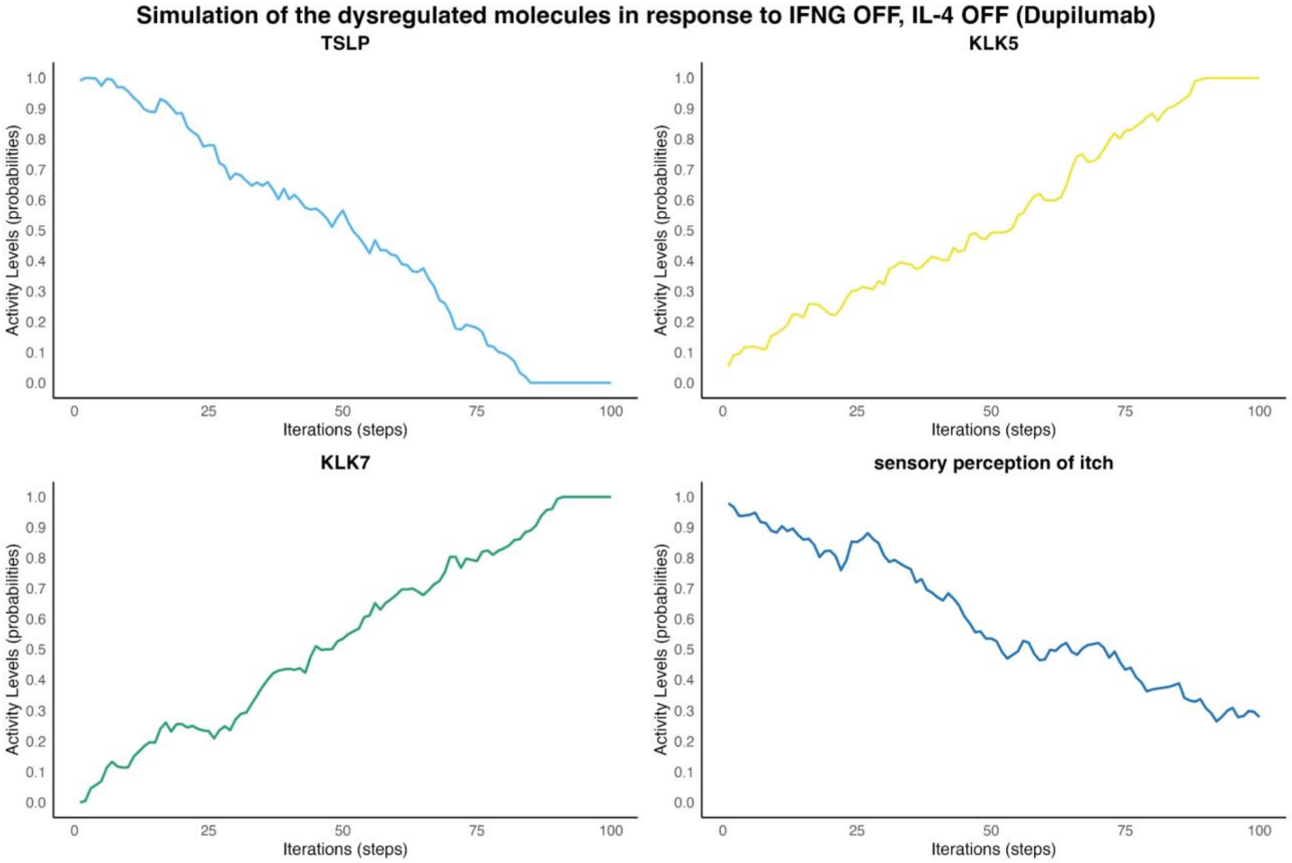


Figure E2. **Boolean network simulation of IFNG effect on sensory itch perception after dupilumab treatment:**. When IFNG is not expressed (IFNG OFF) in the presence of dupilumab (IL4 OFF), we observe a decrease in TSLP levels, an increase in KLK5 and KLK7 levels and a reduction of around 70% in the activity level of sensory perception of itch.
